# Histidine and its uptake are essential for the growth of *Staphylococcus aureus* at low pH

**DOI:** 10.1101/2023.07.25.550546

**Authors:** Catrin M. Beetham, Christopher F. Schuster, Marina Santiago, Suzanne Walker, Angelika Gründling

## Abstract

*Staphylococcus aureus* is an opportunistic pathogen capable of causing many different human diseases. During colonization and infection, *S. aureus* will encounter a range of hostile environments, including acidic conditions such as those found on the skin and within macrophages. However, little is known about the mechanisms that *S. aureus* uses to detect and respond to low pH. Here, we employed a transposon sequencing approach to determine on a genome-wide level the genes required or detrimental for growth at low pH. We identified 31 genes that were essential for the growth of *S. aureus* at pH 4.5 and confirmed the importance of many of them through follow up experiments using mutant strains inactivated for individual genes. Most of the genes identified code for proteins with functions in cell wall assembly and maintenance. These data suggest that the cell wall has a more important role than previously appreciated in promoting bacterial survival when under acid stress. We also identified several novel processes previously not linked to the acid stress response in *S. aureus*. These include aerobic respiration and histidine transport, the latter by showing that one of the most important genes, *SAUSA300_0846*, codes for a previously uncharacterized histidine transporter. We show that an *S. aureus SAUSA300_0846* mutant is unable to maintain its cytosolic pH, thereby revealing an important function for histidine and its transport in buffering the intracellular pH in bacteria.

**Author summary:** *Staphylococcus aureus* is an important human bacterial pathogen that can cause a range of diseases. During infection, the pathogen will encounter a hostile environment within the human host, including acidic conditions such as those found on the skin and within macrophages. The bacterium has developed sophisticated strategies to survive and grow under such harsh conditions. Here we performed a genome wide screen to identify factors that are required by this pathogen to survive under acid stress conditions and identified several novel processes including histidine uptake. Understanding the response of *S. aureus* to deal with acid stress conditions will help us better manage infections.

## Introduction

*Staphylococcus aureus* is a Gram-positive commensal bacterium that is found on the skin, in the respiratory tract, and in the nasal passage (1). However, *S. aureus* also causes a variety of infections ranging from skin and soft tissue infections to bacteraemia (2). To cause such a wide range of infections, *S. aureus* needs to be able to survive in different niches in the host, many of which are inhospitable to bacterial growth. One such environment is low pH, which is present on the skin, the stomach, and in phagosomes during macrophage uptake.

Low pH can adversely affect bacteria in several ways. Firstly, the increased extracellular proton concentration affects the electrochemical gradient of protons across the membrane, known as the proton motive force (PMF). Bacteria use the PMF to generate ATP, and changing the pH has been shown to affect ATP synthesis (3). Protons can also cause damage inside the cell by increasing formation of reactive oxygen species (ROS) (4, 5), affecting the activity of enzymes that have an optimal pH range, causing denaturation and misfolding of proteins due to different charges on amino acids and promoting DNA damage due to depurination (6). Therefore, bacteria have developed a range of mechanisms to deal with low pH (reviewed in 7, reviewed in 8). Firstly, bacteria can reduce the permeability of their cell membrane and cell wall to protons by changing either its composition or charge. Notably in Gram-positive bacteria, the addition of the positively charged D-ala via the *dlt* operon onto the teichoic acids has been shown to be important for growth and survival at low pH (9, 10). Secondly, bacteria can pump excess protons out of the cell via proton pumps such as the F_0_F_1_-ATPase (11). Thirdly, bacteria can increase the rate of reactions which consume protons, such as amino acid decarboxylation. Common pathways include glutamate, lysine, or arginine decarboxylation, with the resulting products and protons being exported from the cell (12). An alternative method to buffer the cytoplasmic pH is to produce alkaline molecules such as ammonia. Ammonia can be produced either by the arginine deaminase system (reviewed in 13), or from urea via the urease enzyme (reviewed in 14). Ammonia is readily protonated to form ammonium, thus consuming a proton. Finally, bacteria can respond to low pH by upregulating processes which protect against pH induced damage, or repair macromolecules following damage. This can include chaperone proteins to prevent pH-induced protein denaturation, proteases to degrade damaged proteins, ROS detoxification systems, or DNA repair processes (reviewed in 7, 15, 16).

While much work has been done in Gram-negative bacteria such as *Escherichia coli* and *Salmonella enterica*, less is known how Gram-positive bacteria, and in particular *S. aureus,* copes with acid stress. Most information on the acid stress response of *S. aureus* has been gained from transcriptional studies (reviewed in 17, 18, 19, 20, 21). Some of the most upregulated genes in these studies were the *ureaA-F* and *nixA* genes, encoding the urease enzyme UreAB and accessory proteins UreDEF, and the high-affinity nickel transporter NixA required for full urease activity. It has also been experimentally verified that the production of ammonia from urea via the urease enzyme is essential for the survival of *S. aureus* under weak acid stress as well as required for pathogenicity of *S. aureus* (22). Furthermore, *S. aureus* can produce ammonia via the arginine deaminase system and the arginine catabolic mobile element (ACME) facilitates this process (23). Other genes that were transcriptionally upregulated encode proteins involved in cell wall assembly and modulation of cell surface charge. These include genes involved in capsular polysaccharide biosynthesis (18), and the *dlt* genes, which encode enzymes that catalyse the addition of positive charges onto the cell surface (20). In several of the transcriptional studies, upregulation of *groEL* and *groES* genes encoding for chaperone proteins or *clpC*, *clpP*, and *clpB* encoding Clp proteases was seen (18, 19, 20, 21). Furthermore, genes involved in DNA repair mechanisms such as *rexAB* (19) and genes involved in ROS detoxification were also upregulated (19). Finally, intracellular levels of the signalling nucleotide cyclic di-adenosine monophosphate (c-di-AMP) have been shown to correlate with acid sensitivity. A low c-di-AMP producing strain had decreased growth at pH 4.5, while a high c-di-AMP producing strain displayed the opposite phenotype (24). The mechanism for this is currently still unknow.

Despite the wealth of information from these transcriptional studies, such studies do not take into account post-transcriptional or post-translational modifications. Secondly, in most of these studies only initial responses within 5-30 minutes of acid shock were investigated. Finally, very few phenotypic studies have been performed to experimentally validate individual pathways involved in the acid stress response of *S. aureus*. Here, we employed a transposon sequencing approach to determine on a whole-genome levels genes that are required or dispensable (meaning their disruption might be beneficial) for growth of *S. aureus* in low pH. The genes highlighted as important for growth in low pH medium support some previously identified *S. aureus* acid stress responses, including the importance of cell surface charge. However, our study suggests that other cell wall factors such as peptidoglycan hydrolases play a key role. Several genes previously not linked to acid stress were also identified, including the terminal oxidase genes *qoxAB* and *SAUSA300_0846*. A further characterisation revealed that SAUSA300_0846 is the main histidine transporter in *S. aureus.* Finally, we show that histidine and its uptake are important for the growth of *S. aureus* under acidic conditions by helping bacteria maintain their intracellular pH.

## Results

### Using Tn-Seq to identify genes required or detrimental for the growth of *S. aureus* under acid stress conditions

To identify genes required or detrimental for the growth of *S. aureus* under acidic stress conditions, Tn-seq experiments were performed using two *S. aureus* transposon mutant libraries in the USA300 TCH1516-derived strain background TM283 (25, 26). These libraries will be referred to as libraries, A and B, and contain approximately 600,000 and > 1 million pooled colonies, respectively. The pools of transposon mutants were grown for 10 generations in standard TSB (pH 7.3), and under acid stress conditions in TSB pH 5.5 or TSB pH 4.5. The bacteria were subsequently harvested, and transposon insertion sites mapped by sequencing (Fig. 1A). Cultures propagated at pH 5.5 grew at a similar rate to the cultures grown at pH 7.3, while those incubated at pH 4.5 exhibited a noticeable reduction in growth rate (Fig. 1B). Library A grown at pH 4.5 did not reach the desired OD_600_ value due to clumping. This culture was harvested at an OD_600_ of 1.0, but at a time point when it was expected to reach an OD_600_ of 1.4. For both libraries and under all growth conditions, transposon insertions were found to be distributed throughout the genome, indicating good coverage (Fig. S1). Next, the number of transposon insertions per gene following growth in the low pH media was compared to the number of insertions when propagated at neutral pH (Table S1-4). Genes with at most half as many insertions when grown at low pH than at neutral pH were classified as essential/required for growth under acid stress conditions. Detrimental or dispensable genes were defined as those with at least twice as many insertions when grown at low pH than at neutral pH. These are genes that when inactivated might provide a growth advantage under acid stress. Volcano plots were prepared by plotting the fold-changes in transposon-insertion numbers per gene against q-values (Fig. 1C and 1D). As expected, a greater number of essential genes were identified under pH 4.5 than pH 5.5. Next, we sought to identify common essential or dispensable genes between the libraries A and B, with the assumption that any genes that appear in both libraries are more likely to be relevant. To further exclude false positive hits, only genes with a Benjamini-Hochberg corrected p-value of ≤ 0.1 were considered for this analysis. Using these cut-offs, we classified 31 genes as essential/required and 10 genes as dispensable/detrimental for growth under pH 4.5 (Fig. 1E, Table 1). Under the pH 5.5 growth conditions, only 5 genes were identified as essential and 15 genes as dispensable (Fig. 1F). Since only very few essential genes were identified under the pH 5.5 condition, an indication that the stress was not sufficient, we focussed our further analysis on the genes identified at pH 4.5.

**Figure 1.**
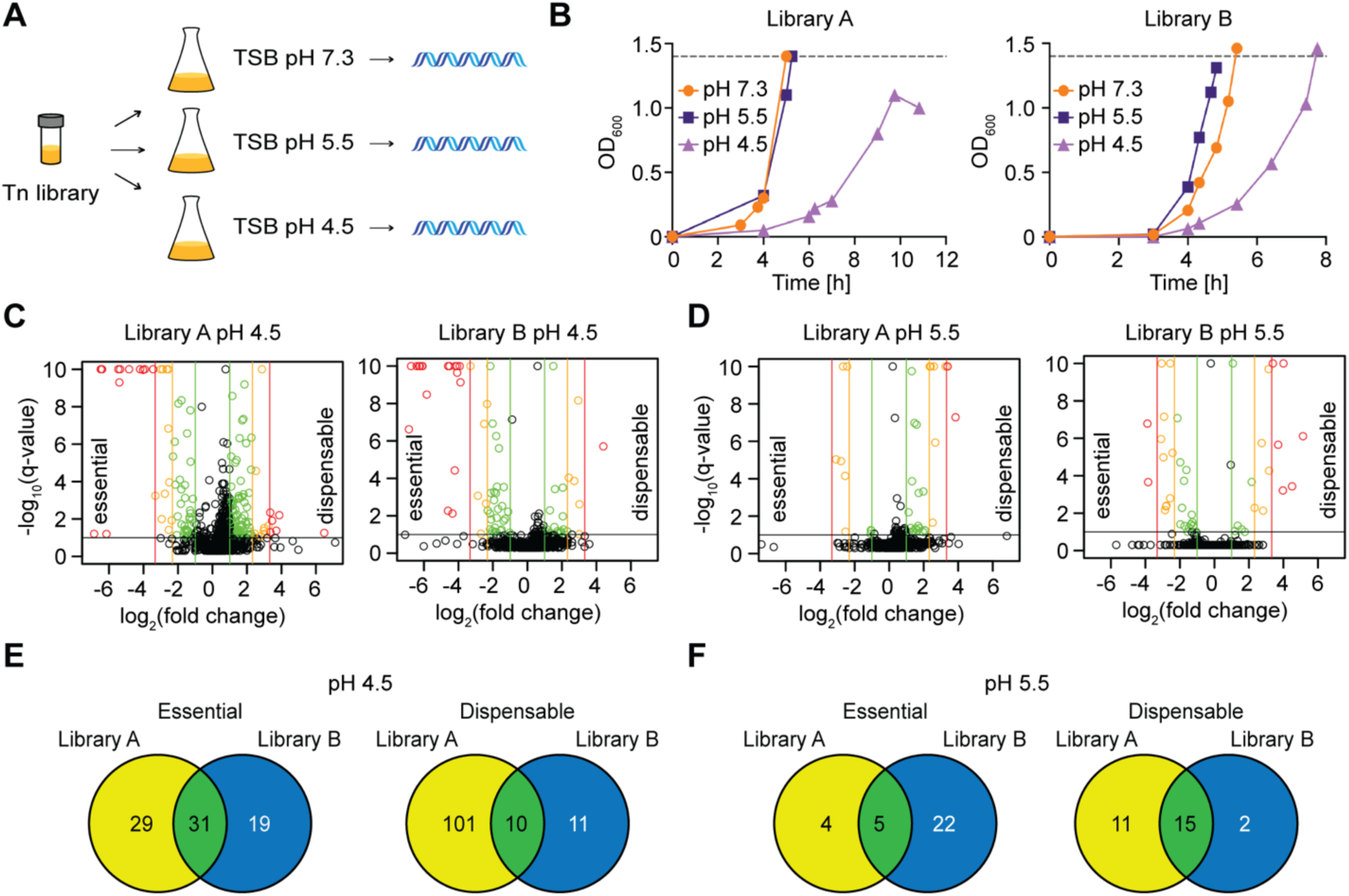
Tn-Seq identifies genes required or detrimental for the growth of *S. aureus* under low pH. (A) Schematic representation of the Tn-Seq experimental protocol. The transposon library was grown in TSB at pH 7.3, pH 5.5, or pH 4.5 for approximately 10 generations, bacteria harvested and transposon insertion sites identified by sequencing. (B) Bacterial growth curves. The growth of two transposon mutant pools (library A and library B) in TSB pH 7.3, pH 5.5, or pH 4.5 medium was monitored by taking OD_600_ readings at defined time points. The dotted line represents the 10-generation cut off point at which the bacteria were harvested. The culture of Library A grown at pH 4.5 did not reach the desired OD_600_ value due to clumping and was instead harvested after 11 h of growth. (C and D) Volcano plots. As a visual representation of the number of essential and dispensable genes obtained following growth of *S. aureus* in (C) TSB pH 4.5 or (D) TSB pH 5.5 compared to pH 7.3 volcano plots were generated. On the x-axis the fold change in the number of transposon insertions per gene was plotted on a log_2_ scale with negative values indicating essential genes and positive values indicating dispensable genes. Vertical green, orange or red lines indicate 2-, 5- or 10-fold changes, respectively. The y-axes are the *q*-values (Benjamini–Hochberg corrected *p*-values) and the black horizontal line indicates a *q*-value of 0.1. Each dot represents one gene and colouring follows the fold-change scheme whenever the *q*-value threshold was met. Very small *q*-values were truncated to fit onto the graph. (E and F) Venn diagrams. The list of genes with a 2-fold decrease (essential genes) or a 2-fold increase (dispensable genes) in the number of transposon insertions and a Benjamini–Hochberg value of ≤ 0.1 were compared between libraries A and B and displayed in Venn diagrams for (E) pH 4.5 and (F) pH 5.5 stress conditions. The overlapping 31 essential and 10 dispensable genes for the pH 4.5 growth condition are listed in Table 1.

**Table 1:** Genes identified by Tn-Seq as essential or dispensable for growth of *S. aureus* in TSB pH 4.5 medium.

| Locus Tag<br>SAUSA300_ | Gene<br>Name | Annotated Gene Function | Phenotype in<br>TnSeq at pH<br>4.5 | Average<br>Ratio | Growth plate assay<br>phenotype and other<br>comments |
| --- | --- | --- | --- | --- | --- |
| **Essential genes: Disruption of these genes should decrease the growth of *S. aureus* at low pH** |  |  |  |  |  |
| **0846** |  | Predicted transporter | Essential | 0.01 | Acid Sensitive |
| **0646** | *graS* | Signal transduction histidine kinase | Essential | 0.02 | Acid Sensitive |
| **1865** | *vraR* | DNA-binding response regulator | Essential | 0.02 | Acid Sensitive |
| **0648** | *vraG* | ABC transporter permease | Essential | 0.02 | Acid Sensitive |
| **2389** |  | MSF family permease | Essential | 0.03 | Acid Sensitive |
| **1593** | *secDF* | Protein translocase | Essential | 0.05 | No mutant strain available in NTML collection |
| **1255** | *mprF* | Oxacillin resistance related FmtC protein | Essential | 0.06 | Acid Sensitive |
| **0429** |  |  | Essential | 0.07 | No phenotype |
| **0543** |  |  | Essential | 0.07 | Acid Sensitive |
| **0959** | *fmtA* | Teichoic acid D-ala esterase | Essential | 0.08 | Acid Sensitive |
| **0482** |  | Polysaccharide biosynthesis protein | Essential | 0.10 | No phenotype |
| **2645** | *gidA* | tRNA uridine-5-carboxymethylaminomethyl synthesis enzyme | Essential | 0.10 | No mutant strain available in NTML collection |
| **0481** |  | Transcription-repair coupling factor | Essential | 0.11 | Acid Sensitive |
| **0836** | *dltB* | D-alanyl-lipoteichoic acid biosynthesis protein | Essential | 0.11 | Acid Sensitive |
| **1866** | *vraS* | Two-component sensor histidine kinase | Essential | 0.12 | Acid Sensitive |
| **0855** | *mnhA1* | Monovalent cation/H<sup>+</sup> antiporter subunit A | Essential | 0.20 | No mutant strain available in NTML collection |
| **2646** | *trmE* | tRNA modification GTPase | Essential | 0.21 | Tn insertion site could not be confirmed in NTML strain |
| **1158** |  |  | Essential | 0.22 | No mutant strain available in NTML collection |
| **0957** |  |  | Essential | 0.22 | No phenotype |
| **1720** | *sagB* |  | Essential | 0.24 | No phenotype |
| **1518** | *cshB* | DEAD-box ATP dependent DNA helicase | Essential | 0.24 | Acid Sensitive |
| **2506** | *isaA* | Immunodominant antigen A | Essential | 0.28 | No mutant strain available in NTML collection |
| **0672** | *mgrA* | MarR family transcription factor | Essential | 0.31 | No mutant strain available in NTML collection |
| **0730** | *gdpS* | GGDEF domain containing protein | Essential | 0.31 | No mutant strain available in NTML collection |
| **2282** | *spdC* |  | Essential | 0.31 | No phenotype |
| **0726** |  |  | Essential | 0.32 | No mutant strain available in NTML collection |
| **1442** | *srrA* | Respiratory response protein | Essential | 0.33 | Acid Sensitive |
| **0962** | *qoxB* | Quinol oxidase, subunit I | Essential | 0.35 | Acid Sensitive |
| **0762** | *secG* | Pre-protein translocase subunit SecG | Essential | 0.36 | No mutant strain available in NTML collection |
| **0963** | *qoxA* | Quinol oxidase, subunit II | Essential | 0.40 | Acid Sensitive |
| **1685** |  |  | Essential | 0.44 | No mutant strain available in NTML collection |
| **Dispensable genes: Disruption of these genes should improve the growth of *S. aureus* at low pH** |  |  |  |  |  |
| **1636** | *polA* | DNA superfamily I polymerase | Dispensable | 2.52 | No phenotype |
| **1544** | *lepA* | GTP-binding protein | Dispensable | 3.21 | No phenotype |
| **0759** | *pgm* | Phosphoglyceromutase | Dispensable | 3.35 | Acid Sensitive |
| **2174** | *ecfT* | Cobalt transport family protein | Dispensable | 3.58 | Tn insertion site could not be confirmed in NTML strain |
| **0746** |  |  | Dispensable | 3.63 | Tn insertion site could not be confirmed in NTML strain |
| **2643** | *noc* | ParB family chromosome partitioning family | Dispensable | 3.66 | No phenotype |
| **2055** | *murA* | UDP-N-acetylglucosamine 1-carboxyvinyltransferase | Dispensable | 4.12 | No phenotype |
| **1043** | *mutS2* | Recombination and DNA strand exchange inhibitor protein | Dispensable | 5.10 | No phenotype |
| **1509** | *actH* | Rhomboid family peptidase | Dispensable | 5.52 | No mutant strain available in NTML collection |
| **1588** | *lytH* | N-acetylmuramoyl-L-alanine amidase | Dispensable | 12.96 | No phenotype |

### Tn-Seq highlights the bacterial cell wall as a key component of the acid stress response of *S. aureus*

To identify the main pathways that are required for *S. aureus* to grow under low pH conditions, we examined the list of genes highlighted in the Tn-seq experiments under the pH 4.5 growth condition in more detail (Table 1). Many genes coding for factors connected to the bacterial cell wall were found to be required under low pH growth conditions (Fig. 2). *vraS* and *vraR* coding for the VraSR two-component system, which detects cell-wall stress, were on the list of essential genes. Several peptidoglycan hydrolases were also on the list, including SpdC and SagB, which form a protein complex required for the release of nascent peptidoglycan during daughter cell formation, and IsaA, the immunodominant staphylococcal antigen A, which is a predicted lytic transglycosylase enzyme (27, 28, 29). Other pathways linked to the cell wall included those involved in modulating cell surface charge. *graS* and *vraG,* coding for two membrane bound sensory components of the GraXRS-VraFG five-component system, were identified. This system detects and responds to cell wall stressors such as cationic antimicrobial peptides, leading to the expression of factors required for the addition of positive charges to the cell surface (30). Several of the *graXRS-vraFG* target genes were also highlighted as essential in the Tn-Seq assay. These include *dltB* from the *dlt* operon, which is required for the addition of positively charged D-alanine residues onto teichoic acids, and *mprF*, which is responsible for the synthesis of the positively charged membrane lipid lysyl-phosphatidylglycerol. Finally, *fmtA*, encoding an enzyme which has recently been shown to be a D-amino acid esterase that removes D-alanines from teichoic acids, was also highlighted as essential under low pH growth conditions. Interestingly *lytH* and *actH,* coding for a peptidoglycan amidase and its partner protein ActH and required for LytH activity (31, 32), were identified as the two top dispensable gene under acid stress conditions, indicating that the activity of this amidase is detrimental under these growth conditions.

**Figure 2:**
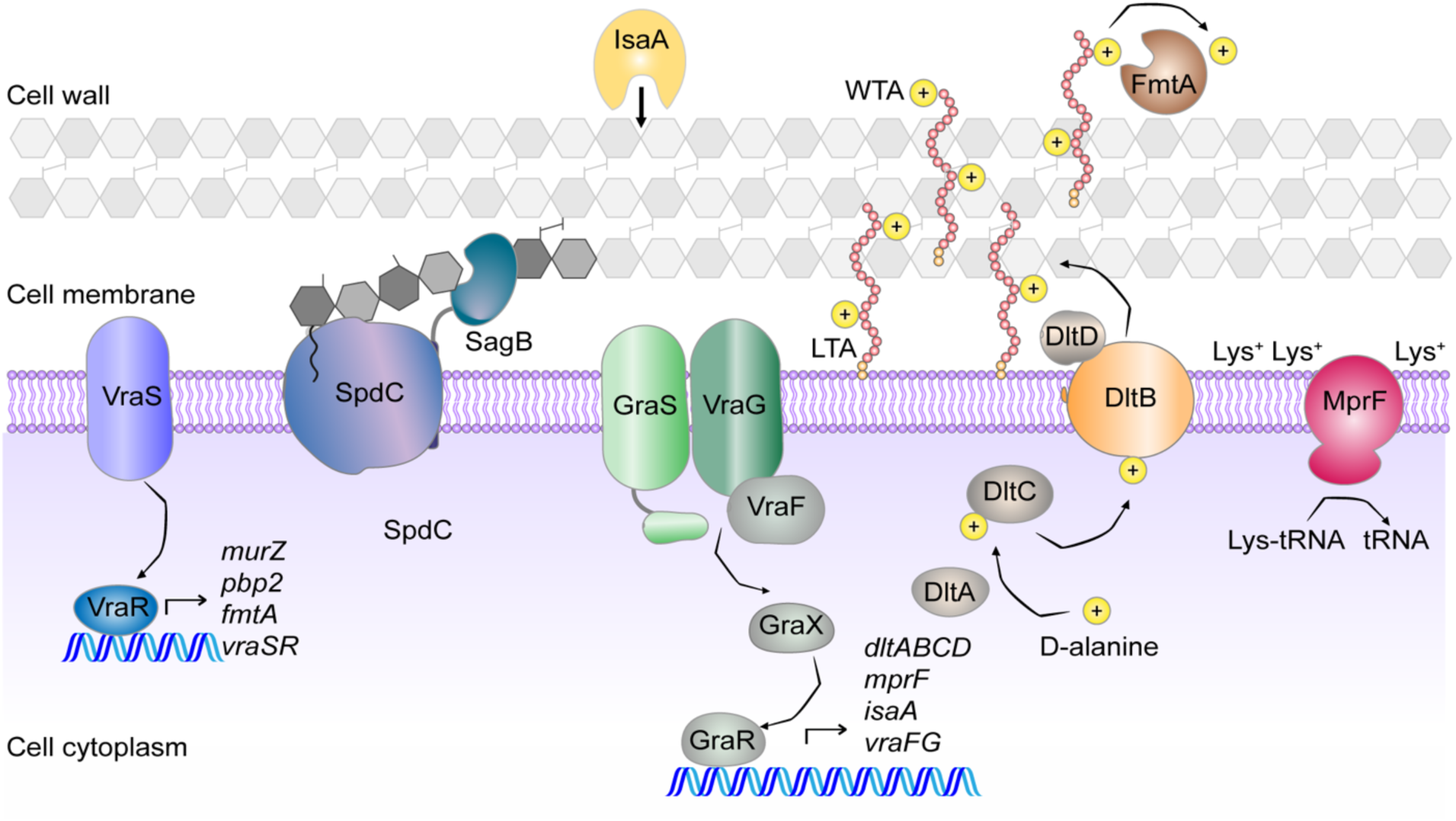
A number of factors involved in the cell wall remodelling were identified as essential in the TnSeq experiment. Schematic representation of the *S. aureus* cell wall with proteins highlighted in this study as essential for growth in TSB pH 4.5 and involved in cell wall synthesis or remodelling depicted. If the proteins are part of complexes, the proteins identified in the Tn-seq experiment are shown in colour while the others are shown in grey.

In addition, there were several pathways highlighted as essential that have not previously been associated with the acidic stress response in *S. aureus*. One such pathway is aerobic respiration. *srrA* was identified, which codes for the response regulator of the SrrAB two-component system that regulates genes involved in aerobic respiration. These genes include *qoxA* and *qoxB*, coding for subunits of the Qox cytochrome *aa*_3_-type quinol oxidase, a proton pumping terminal oxidase present in *S. aureus*. Consistent with this, the *qoxA* and *qoxB* genes were also highlighted as required for growth under low pH growth conditions in the Tn-Seq experiment. Other genes that were identified were *SAUSA300_0846* and *mhnA1* coding for predicted cation:proton antiporters, *secDF* coding for protein translocase subunits, and *gidA* coding for a tRNA uridine 5-carboxymethylaminomethyl modification enzyme. Furthermore, several genes with still uncharacterized functions were identified, such as *SAUSA300_2389* coding for a putative MSF family permease, *SAUSA300_0429* coding a predicted phospholipid phosphatase and *SAUSA300_0543* coding for a proposed t-RNA adenosine deaminase. Taken together, these data confirm the robustness of using Tn-Seq as a method to explore the acid stress response of *S. aureus*, since previously known pathways were identified as important for growth in low pH, including the cell wall and cell surface charge (20, 33, 34, 35). In addition, several novel pathways and genes were identified in our experiments such as aerobic respiration and *SAUSA300_0846* coding for a predicted transporter.

### Confirmation of genes required for the growth of *S. aureus* under low pH

To further explore to what extent each gene identified in the Tn-Seq assay contributes to the ability of *S. aureus* to grow under low pH conditions, we investigated the growth of individual *S. aureus* mutant strains. To do this, we used defined transposon mutants available from the Nebraska Mutant Transposon Library (NTML) (36), or mutant strains available from other collections. However not all identified genes could be further investigated (Table 1) since we were unable to confirm the transposon insertion site in some of the NMTL strains and some genes are thought to be essential for normal growth and thus no mutant strains existed in our collection. In the end, we assayed the growth of 20 mutant strains from the list of 31 essential genes and 7 mutant strains from the list of 10 dispensable genes. The growth of each mutant was tested by spotting serial dilutions of overnight cultures onto TSA pH 7.3 or TSA pH 4.5 plates. None of the strains displayed any drastic difference in the number of colony forming units (CFUs) on TSA pH 7.3, showing that inactivating these genes does not confer any severe growth defect under neutral pH conditions (Fig. S2). On the other hand, several of the mutant strains displayed a growth defect on TSA pH 4.5 plates (Fig. 3). Most of the mutant strains carrying transposon insertions or deletions in cell wall related genes showed growth defects on the TSA pH 4.5 plates. This included *graS*, *vraG*, *vraR*, *vraS*, *mprF*, *fmtA*, and *dltD* mutants (Fig. 3A-C), thus confirming the importance of the cell wall in the acid stress response of *S. aureus* (Fig. 2). Inactivating genes involved in aerobic respiration also resulted in a reduced ability of the strains to grow on TSA pH 4.5 plates. The *srrA* mutant showed a severe phenotype, and a slight growth reduction was also observed for *qoxA* and *qoxB* mutants (Fig. 3E). Other strains which showed a reduced ability to grow on pH 4.5 plates included mutants with transposon insertions in *SAUSA300_0846* (predicted transporter), *SAUSA300_2389* (predicted MFS permease), *SAUSA300_0543* (predicted t-RNA adenosine deaminase), and a slight reduction in growth was seen for *SAUSA300_0481* (transcription-repair coupling factor Mfd) and *SAUSA300_1518* (DEAD-box ATP dependent DNA helicase CshB) mutants (Fig. 3F, G). However, not all the mutant strains displayed a reduced ability to grow on TSA pH 4.5, despite being identified as essential in the Tn-Seq experiment. These included *SAUSA300_0482*, *SAUSA300_0957*, *sagB*, *spdC*, and *SAUSA300_0429* mutants (Fig. 3D,F,G).

**Figure 3:**
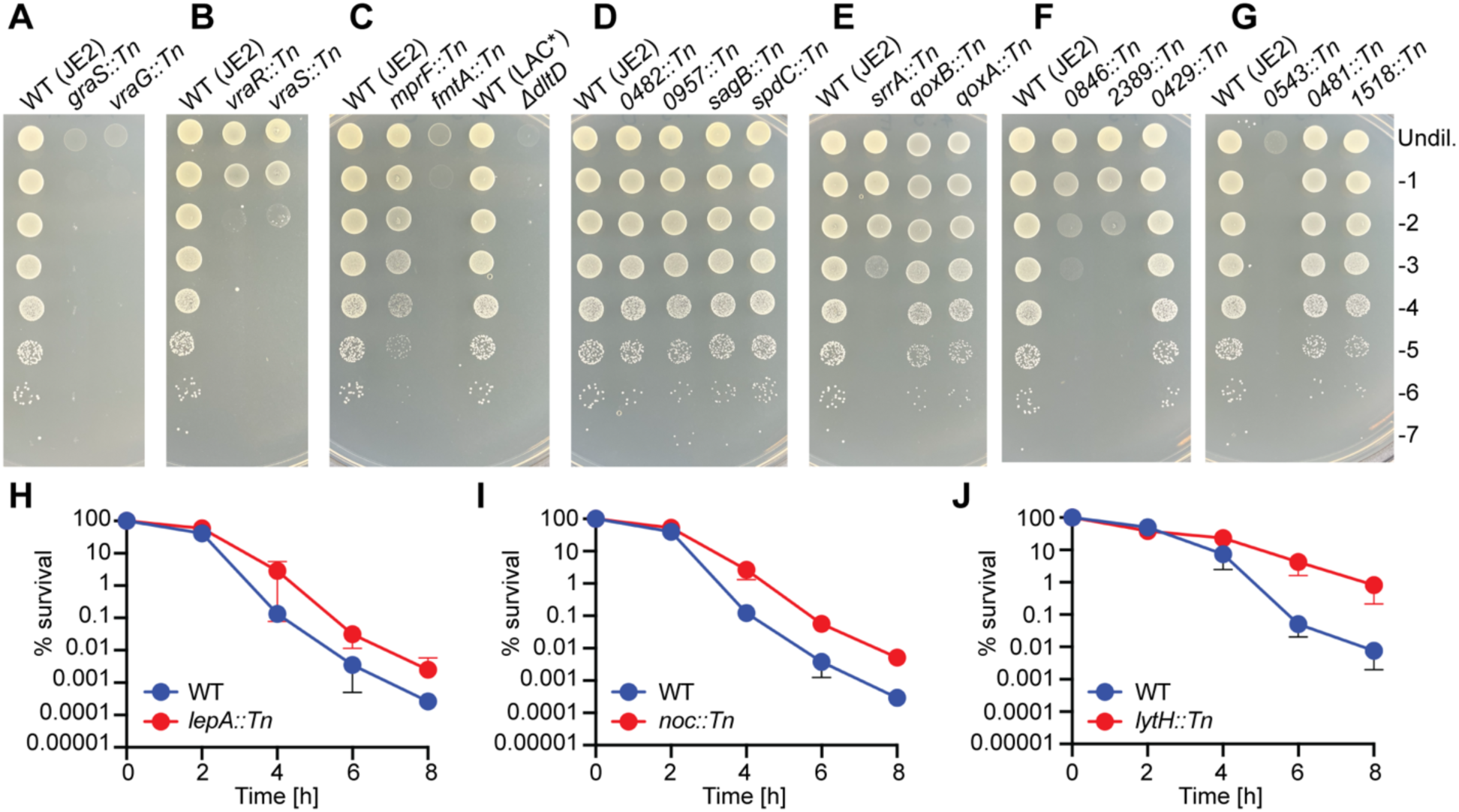
Growth plate and survival analysis of *S. aureus* mutant strains with transposon insertions in genes identified as essential or dispensable for growth at pH 4.5. (A-G) Bacterial growth on TSA pH 4.5 plates. Overnight cultures of the indicated WT and mutant strains were serially diluted and aliquots spotted onto TSA pH 4.5 plates. Images were taken following 24 h incubation at 37 °C. Each image is a representative of three experiments. (H-J) Acid stress survival curves. Overnight cultures of WT JE2 as well as (H) JE2 *lepA::Tn*, (I) JE2 *noc::Tn* or (J) JE2 *lytH::Tn* were washed and diluted into TSB pH 2.5 medium. Immediately afterwards (T = 0 h) and at 2 h intervals up to 8 h aliquots were removed and CFUs determined by plating dilutions onto TSA plates. The CFU count at T = 0 h was set to 100% for each strain and % survival at the subsequent time points calculated. The average and standard deviations of the % survival from three independent experiments were plotted.

None of the 7 strains, which were available to us, with transposon insertions in the dispensable genes displayed an increased growth on TSA pH 4.5 (Fig. S3). We hypothesized that this may be because the assay is not stringent enough to detect an increase in growth at pH 4.5 due to the robust growth of the WT strain under this condition. We therefore also assayed the survival of the WT and dispensable mutants in liquid medium at a much lower pH of 2.5. In this assay, three of the strains had increased survival compared to the WT in TSB pH 2.5 (Fig 3H, I, J). These were *lepA::Tn*, *noc::Tn*, and *lytH::Tn*. The latter, which had the highest ratio in the Tn-Seq assay of 12.96, also displayed the greatest increase in survival, with a 100-fold increase in CFU ml^-1^ counts compared to the WT following 8 h incubation in TSB pH 2.5 (Fig 3H, I, J). Overall, of the 20 essential genes investigated, 15 could be confirmed to be important for the growth of *S. aureus* under low pH conditions. Furthermore, for 3 of the 7 dispensable genes investigated it could be shown that their inactivation leads to increased survival of *S. aureus* under low pH conditions. This highlights that Tn-seq is a reliable method for identifying genes required for the growth of bacteria under acid stress conditions. The importance of several novel genes was confirmed including for *SAUSA300_0846* coding for an uncharacterized membrane transporter, which we investigated further as part of this study.

### 0846 is a main histidine transporter in *S. aureus*

*SAUSA300_0846* (hereon called *0846*) was chosen for further study as it had the lowest ratio in the Tn-Seq experiment at pH 4.5, and hence can be deemed as one of the most important factors during low pH conditions. The gene was also identified as essential in the Tn-Seq experiment at pH 5.5. Interestingly, several previous studies have demonstrated that a *0846* mutant displays reduced virulence in mice (37, 38). However, the cellular function of 0846 is unknown, thus potentially highlighting a novel pathway required for growth of bacteria under low pH conditions and during infection. To confirm that the acid sensitive growth phenotype of the *0846::Tn* mutant strain is due to the disruption of the *0846* gene, the *0846::Tn* region was transferred to a clean *S. aureus* LAC* background strain. In addition, a complementation strain was generated by introducing the single site integration plasmid pCL55-*0846* into the LAC* *0846::Tn* mutant strain for expression of *0846* under its native promoter control. Strain LAC* *0846::Tn* containing the empty pCL55 plasmid showed the expected growth defect at low pH, both on TSA pH 4.5 agar plates as well as in liquid medium (Fig. 4A, B). The growth was restored to WT levels in the complementation strain LAC* *0846::Tn* pCL55-*0846*, confirming that the acid sensitivity growth phenotype is due to disruption of *0846* (Fig. 4).

**Figure 4:**
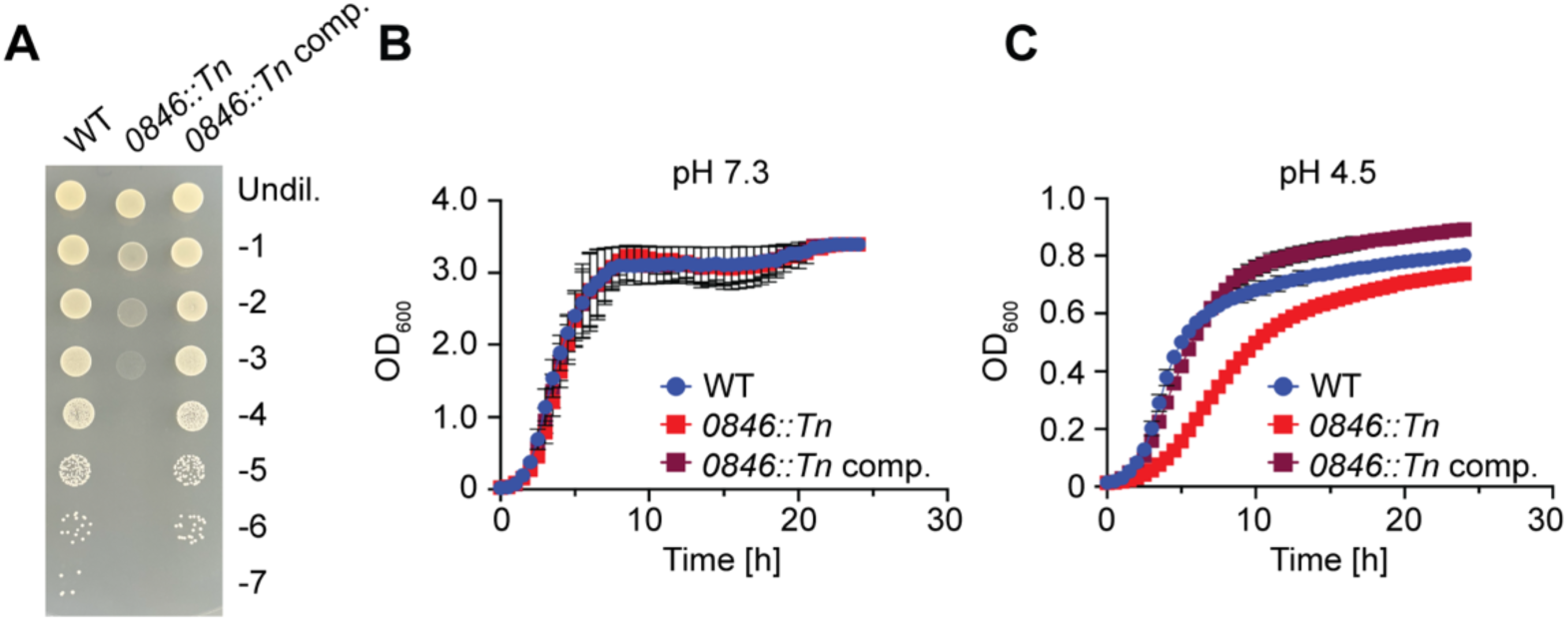
Genetic complementation restores the acid growth ability of the *0846::Tn* mutant strain. (A) Bacterial growth on TSA pH 4.5 plates. Overnight cultures of LAC* pCL55 (WT) LAC* *0846::Tn* pCL55 (*0846::Tn*), and the complementation strain LAC* *0846::Tn* pCL55-*0846* (*0846::Tn* comp.) were serially diluted and aliquots spotted on TSA pH 4.5. Images were taken following 24 h incubation at 37 °C. Each image is representative of three experiments. (B-C) Bacterial growth curves. The same strains as in (A) were grown in (B) TSB pH 7.3 or (C) TSB pH 4.5 medium in 96-well plates and OD_600_ measurements taken at the indicated time points. The average readings and standard deviations of three independent repeats were plotted.

*0846* is part of a two gene *0846-0847* operon and predicted to code for a multi-membrane spanning protein. A literature and bioinformatics analysis revealed two potential functions for 0846. Firstly, it has been annotated as a cation:proton antiporter (CPA) belonging to the NhaC-type of transporter. CPAs have a well-established function during alkaline stress growth conditions in *S. aureus* (39, reviewed in 40, 41). Besides 0846, *S. aureus* has 7 additional potential CPAs; three Cpa1 family transporters, one Cpa2 type transporter, two Cpa3-type Mnh transporters, and two additional NhaC family transporters (39, reviewed in 40, 41). *mnhA1* was identified as essential in the Tn-Seq experiment at pH 4.5 (Table 1), and in a previous study it was shown that *mnhA1* and *cpa1-1* expression increased at pH 6.0, although *0846* did not (41). We therefore hypothesized that, as well as being important under alkaline stress conditions, CPA activity might also be important for the acidic stress response. To explore this further, we investigated the requirement of the different *S. aureus* CPA transporters for growth under low pH conditions. Mutants with transposon insertions in genes coding for these transporters were assayed for their growth ability on TSA pH 4.5 plates. The Mnh transporters however could not be assayed since the *mnh1* operon is essential in LAC, LAC* and JE2-derived strains due to a non-functional *mnh2* operon (36, 39). Only the *0846::tn* mutant, but none of the other five mutant strains with transposon insertions in genes coding for CPAs, displayed a reduced ability to grow on TSA pH 4.5 plates (Fig. 5A). While it is well established that CPA transporter activity is important under alkaline stress conditions, our data suggest that it might not be required for the growth of *S. aureus* under low pH condition, indicating that the importance of 0846 during acid stress might not be due to cation proton antiporter activity.

**Figure 5:**
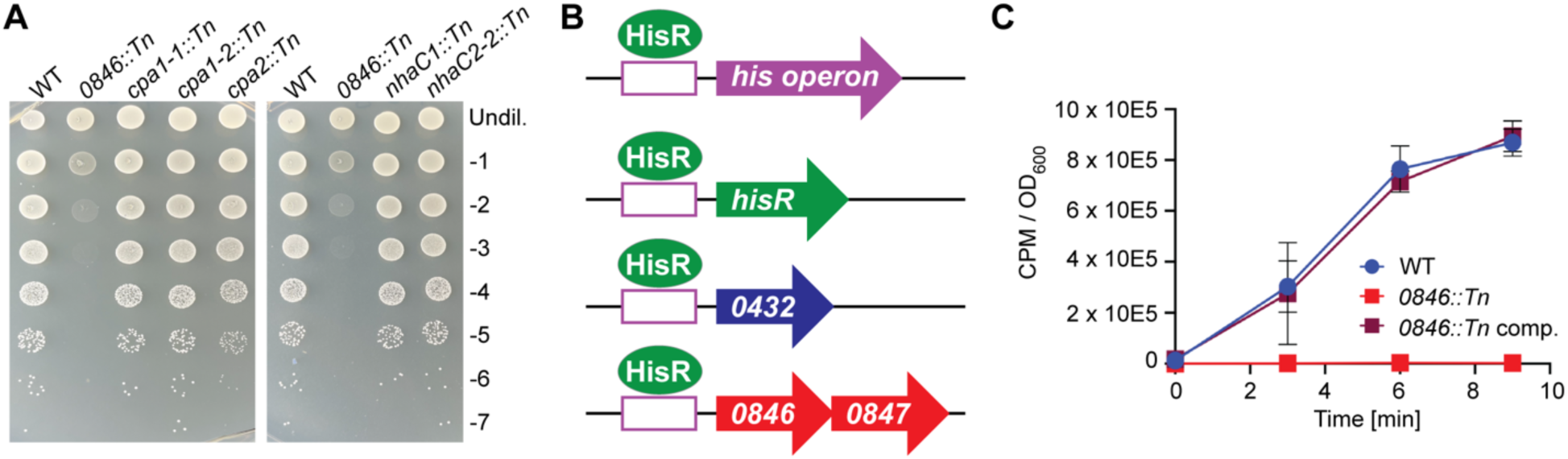
0846 is a main histidine transporter in *S. aureus*. (A) Bacterial growth on TSA pH 4.5 plates. Overnight cultures of JE2 (WT), *0846::Tn*, *cpa1-1::Tn*, *cpa1-2::Tn*, *cpa2::Tn*, *nhaC1::Tn*, *nhaC2::Tn* strains were serially diluted and spotted on TSA pH 4.5 plates. Images were taken following 24 h incubation at 37 °C. Each image is representative of three biological experiments. (B) Schematic representation of the *S. aureus* genes and operons suggested to be regulated by the proposed histidine transcription factor HisR. (C). Histidine uptake assay. ^3^H radiolabelled L-histidine was added to washed mid-log phase cultures of LAC* pCL55 (WT), LAC* *0846::Tn* pCL55 (*0846::Tn*), and the complementation strain LAC* *0846::Tn* pCL55-*0846* (*0846::Tn* comp.). At the indicated time points, culture aliquots were removed, filtered, washed and the accumulated radioactivity in each sample measured as counts per minute (CPM) using a scintillation counter. The CPM values were normalised to OD_600_ values, and the average CPM / OD_600_ and standard deviations of three independent experiments were plotted.

The second annotation for 0846 is as potential YuiF-type histidine permease. Such a function is consistent with other published computational predictions. In a previous bioinformatics study, *S. aureus* operons with predicted upstream transcription factor binding site were identified, including the histidine biosynthesis operon (42, 43) (Fig. 5B). A similar sequence was also identified upstream of a gene since renamed *hisR* (for histidine regulator) and predicted to code for the transcription factor regulating the expression of the histidine biosynthesis operon. This sequence was also found upstream of the *0846-0847* operon (43). To test whether *0846* encodes a histidine transporter, we measured the uptake of radiolabelled histidine in WT *S. aureus* strain LAC*, the isogenic LAC\**0846::Tn* mutant strain (both containing the empty pCL55 plasmid) and the complementation strain LAC\**0846::Tn* pCL55-*0846*. While the WT strain was able to take up histidine, uptake of this amino acid was almost completely eliminated in the *0846::Tn* mutant, and restored again to WT-levels in the complementation strain (Fig. 5C). These data show that under our assay conditions, 0846 functions as the main histidine transporter in *S. aureus*.

### Histidine is important for the growth of *S. aureus* under low pH conditions

To confirm the histidine transport activity of 0846 during bacterial growth, we performed growth curves in chemically defined medium (CDM) with or without 130 mM histidine. In the presence of exogenous histidine, the WT enters exponential growth after approximately 2 h (Fig. 6A). However, when histidine is not present, the WT displays an additional lag in growth of approximately 5 hours. This indicates that *S. aureus* usually takes up histidine to support growth, and if removed it takes a few hours for the bacteria to adapt and synthesize histidine. On the other hand, the *0846::Tn* strain showed a very similar growth phenotype as the wild-type strain in the absence of histidine, regardless of whether or not this amino acid is present (Fig. 6B). This is likely because the *0846::Tn* cannot use exogenous histidine. The complemented *0846::Tn* strain was again able to initiate growth after around 2 h in the presence of histidine (Fig. 6C). Taken together these data are consistent with the notion that *0846* encodes for the main histidine transporter in *S. aureus*.

**Figure 6:**
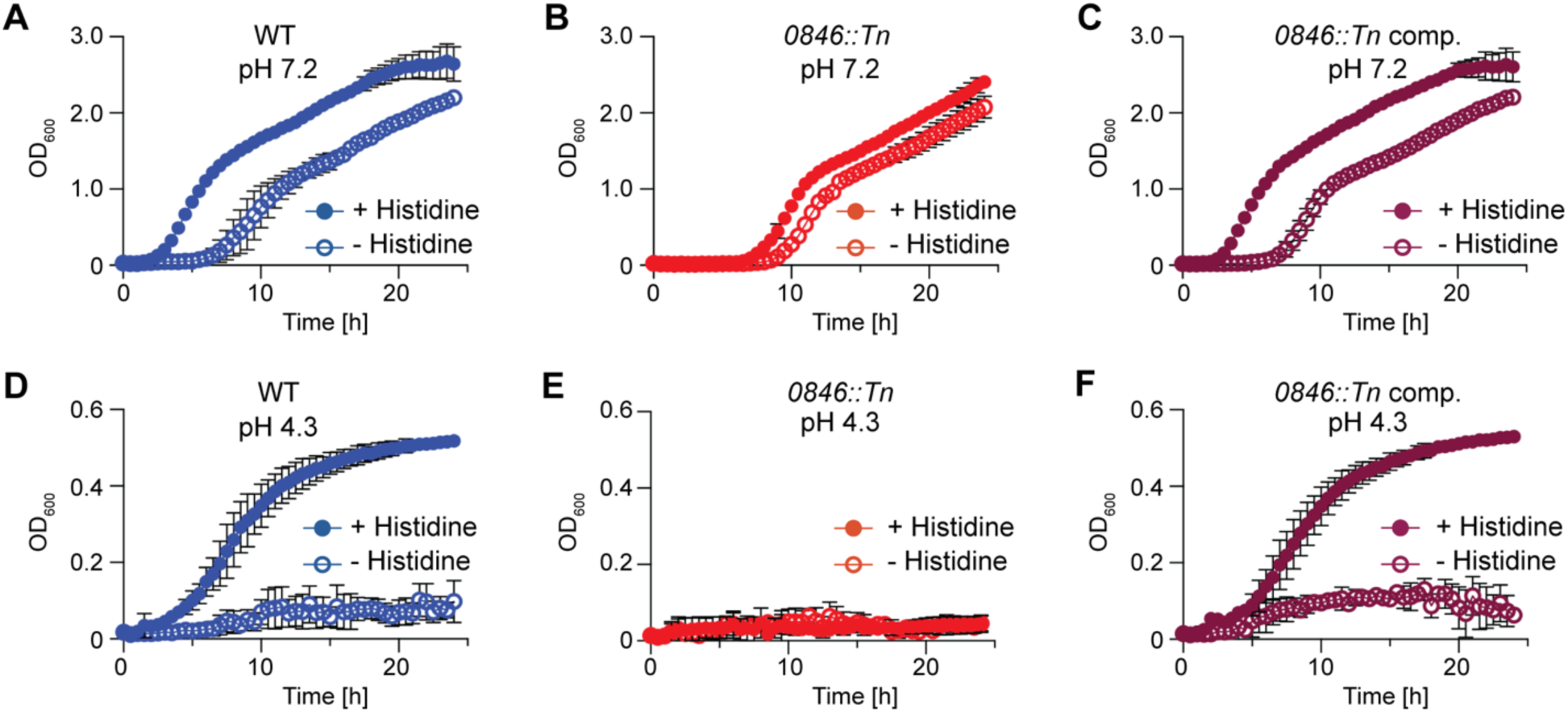
Histidine and its uptake is important for the growth of *S. aureus* under acid stress conditions. (A-C). Bacterial growth curves in CDM pH 7.2. *S. aureus* strains (A) LAC* pCL55 (WT), (B) LAC* *0846::Tn* pCL55 (*0846::Tn*), and (C) LAC* *0846::Tn* pCL55-*0846* (*0846::Tn* comp.) were grown in CDM pH 7.2 with or without 130 mM histidine and OD_600_ readings taken at timed intervals. The average readings and standard deviation of three independent repeats were plotted. (D-F) Bacterial growth curves in CDM pH 4.3. The growth of (D) LAC* pCL55 (WT), (E) LAC* *0846::Tn* pCL55 (*0846::Tn*), and (F) LAC* *0846::Tn* pCL55-*0846* (*0846::Tn* comp.) was monitored and plotted as described for panels A-C, but using CDM pH 4.3 with or without 130 mM histidine.

To assess whether histidine is important for the acid stress response of *S. aureus*, we also compared the growth of the WT, *0846::Tn*, and complemented strain at very stringent pH 4.3 conditions. While the growth of the WT strain was reduced when grown in CDM at pH 4.3 (Fig. 6D) compared to when grown at pH 7.2 (Fig. 6A) it was almost completely unable to grow in the absence of histidine (Fig. 6D). The *0846::Tn* strain, which is unable to take up histidine and was unable to grow at pH 4.3 regardless of whether histidine was present or not (Fig. 6E). Expression of *0846* in the complementation strain restored the ability of the mutant to grow in the presence of histidine at low pH (Fig. 6F). These data indicate that histidine transport via the 0846 transporter is required for the growth of *S. aureus* at low pH.

### The acid sensitivity of the *0846::Tn* mutant strain can be bypassed by inducing the stringent response

To further investigate why *0846* is important for the growth of *S. aureus* under acid stress, we selected for LAC\**0846::Tn* suppressor strains which showed improved growth on TSA pH 4.5 plates. Such strains could be readily obtained (Fig. 7A) and the genomic alterations leading to the improved growth of eight of these suppressor strains were determined by whole genome sequencing (Table 2). Six of the suppressor strains had single nucleotide polymorphisms (SNPs) in *codY* and two in *relA* (Table 2). CodY is a master gene regulator and together with RelA an important component of the stringent response system in *S. aureus* (Fig. 7B). Under nutrient limiting conditions, RelA produces (p)ppGpp from GTP, leading to a reduction of the intracellular concentration of GTP (reviewed in 44). Amongst others, this shifts CodY into a GTP free state, resulting in the transcription of genes involved in nutrient and amino acid biosynthesis. Two of the suppressor strains had the same SNP in *codY* resulting in an early stop codon and likely inactivation of CodY. This suggests that the stringent response is activated in the LAC* *0846::Tn* suppressor strains with mutations in *codY*. The functional consequence of the mutations in *relA* are not as easy to predict. This is because RelA has a (p)ppGpp hydrolase, a (p)ppGpp synthetase and additional C-terminal regulatory domains (45). One of the suppressor strains had an SNP in the hydrolase domain, and the other strain in one of the C-terminal regulatory domains. But given the type of mutations observed in *codY,* we hypothesize that the mutations in *relA* also lead to an activation of the stringent response. We further confirmed the improved growth of two of the *codY* suppressors and the *relA* suppressors under acid stress conditions by performing growth curves in TSB pH 4.5 liquid medium. As a control for these experiments, we included a LAC* *codY::Tn* mutant strain. All suppressor strains behaved similar to the *codY::Tn* mutant and showed improved growth at pH 4.5 compared to the original *0846::Tn* mutant strain (Fig 7C). Taken together, these data suggest that the suppressor strains compensate for the acid-sensitive growth phenotype of the *0846::Tn* mutant by activating the stringent response. These strains likely shift to a state where they synthesize amino acids, which is consistent with the *0846::tn* mutant strain having a defect in amino acid uptake, which is bypassed in the suppressor strains by increasing the synthesis of the amino acid.

**Figure 7:**
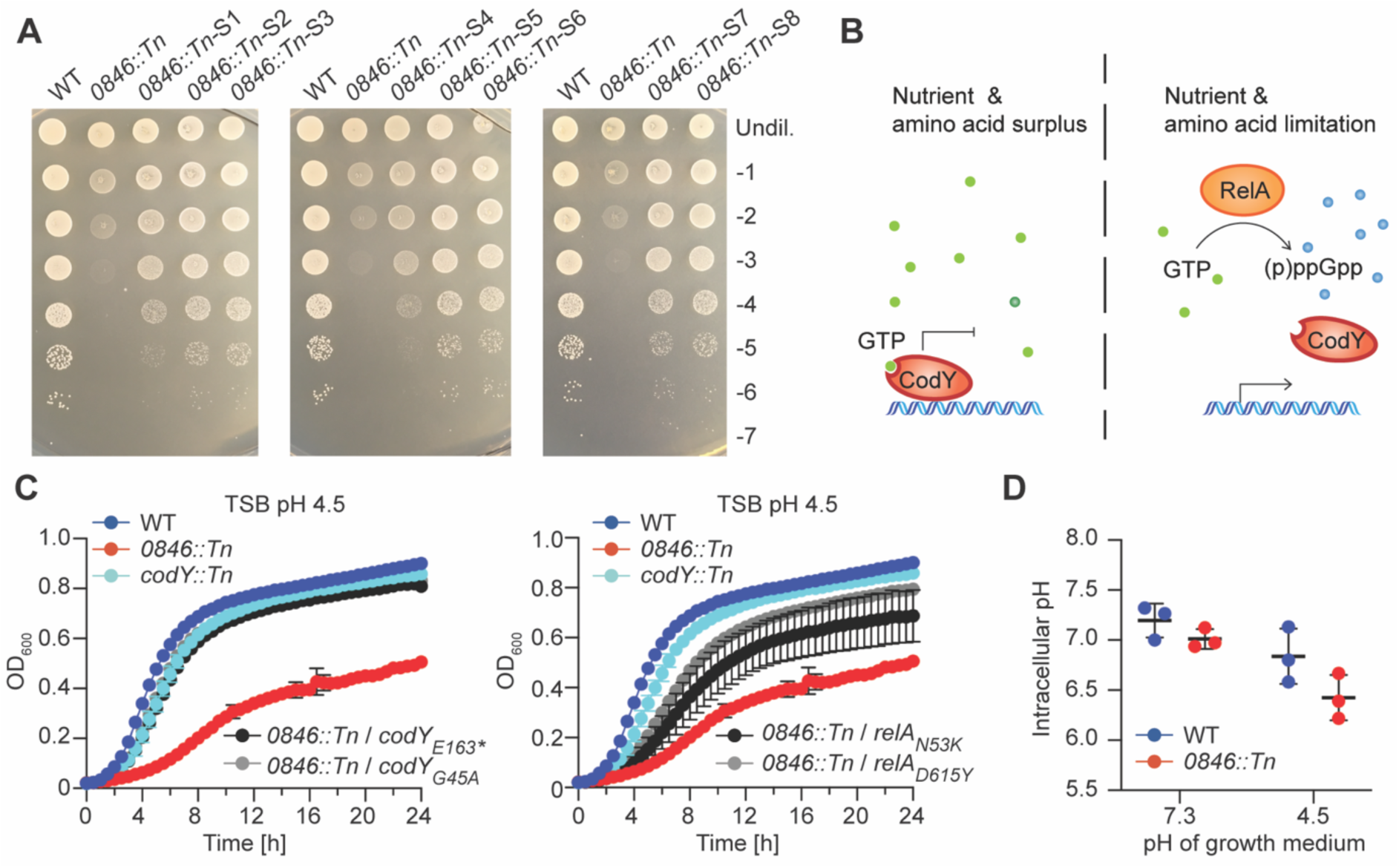
The acid-sensitive phenotype of the *0846::Tn* mutant strain, which has a defect in maintaining the cytosolic pH, can be bypassed by activating the stringent response. (A) Bacterial growth on TSA pH 4.5 plates. Overnight cultures of LAC* (WT), the *0846::Tn* mutant or the suppressor strains *0846::Tn-S1*, *0846::Tn-S2*, *0846::Tn-S3*, *0846::Tn-S4*, *0846::Tn-S5*, *0846::Tn-S6*, *0846::Tn-S7*, *0846::Tn-S8* were serially diluted and aliquots spotted on TSA pH 4.5 plates. Images were taken following 24 h incubation at 37 °C. This experiment was only performed once, but the presence of genomic mutations was confirmed by whole genome sequencing (B). Schematic representation of the stringent response. Under nutrient and amino acid surplus, CodY is in the GTP-bound form, interacts with DNA and prevents gene expression. Under conditions of nutrient or amino acid limitation, RelA produces (p)ppGpp from GTP, decreasing intracellular concentrations of GTP. At low cellular GTP levels, CodY will be in a GTP-unbound state and will no longer bind to DNA, thus allowing expression of the stringent response genes. (C) Bacterial growth curves. LAC* (WT), LAC* *codY::tn, LAC* 0846::tn* mutant and the indicated *LAC* 0846::tn* suppressor strains were grown in (C) TSB pH 4.5 medium and OD_600_ readings taken at timed intervals. The average readings and standard deviations of three independent repeats were plotted. (D) Intracellular pH assay. *S. aureus* strains LAC* (WT) and LAC* *0846::Tn* were grown in TSB pH 7.3 or TSB pH 4.5 to mid-log phase and the intracellular pH determined using the pHrodo Red dye as described in the materials and method section. The average values and standard deviations of three independent experiments were plotted.

**Table 2:** Genomic alterations identified in the LAC\**0846::Tn* suppressor strains.

| Strain number and name | Gene<sup>1</sup> | Mutation<sup>2</sup> | Amino acid change<sup>3</sup> | Frequency<sup>4</sup> |
| --- | --- | --- | --- | --- |
| **6080**<br>**(LAC\* 0846-S1)** | *relA* | SNP C-A | D615Y | 100 |
| **6085**<br>**(LAC\* 0846-S2)** | *codY* | SNP G-T | E163* | 100 |
| **6087**<br>**(LAC\* 0846-S3)** | *codY* | SNP G-T | E163* | 100 |
|  | *SAUSA300_1684* | SNP T-C |  | 70.97 |
| **6090**<br>**(LAC\* 0846-S4)** | *relA* | SNP G-T | N53K | 100 |
| **6099**<br>**(LAC\* 0846-S5)** | *codY* | SNP C-T | L194F | 100 |
| **6111**<br>**(LAC\* 0846-S6)** | *codY* | SNP G-C | G45A | 100 |
| **6113**<br>**(LAC\* 0846-S7)** | *codY* | SNP C-A | R156S | 100 |
| **6118**<br>**(LAC\* 0846-S8)** | *codY* | SNP G-A | G46D | 100 |

<sup>1</sup>Annotated gene where the mutation occurred. <sup>2</sup>Type of mutation with SNP indicating a single nucleotide polymorphism. Base change for SNP indicated by base in reference genome – base in the same position in the suppressor strain. <sup>3</sup>Amino acid change indicating where applicable the initial amino acid, position of amino acid in the encoded protein, and the resulting amino acid following the mutation. AA\* denotes an early stop codon. <sup>4</sup>Frequency indicates the frequency that the mutation was found in the suppressor strain in the genome sequence analysis.

### Histidine uptake is important for the cytoplasmic buffering capability of *S. aureus*

Histidine has a pKa value of around 6.0 and thus can act as a buffer at physiological conditions. To test whether the acid-sensitive phenotype of the *0846::Tn* mutant strain was due to its inability to buffer the cytoplasmic pH, we measured the intracellular pH of WT and the *0846::Tn* mutant strain using the pH sensitive dye pHrodo and a previously published method (38). As expected, the intracellular pH of the WT strain dropped from around 7.2 to 6.8 when the extracellular pH of the growth medium was reduced from 7.3 to 4.5 (Fig. 7D). The intracellular pH of the *0846::Tn* mutant strain during growth in neutral pH medium was already slightly more acidic than that of the WT strain, with a value of around 7.0. This difference in intracellular pH was even more pronounced when the strain was grown in low pH medium, where the intracellular pH dropped to around 6.4. These findings show that the *0846::Tn* strain is unable to regulate its intracellular pH, especially under low pH conditions. This establishes histidine and its transport as new pathway required the survival of *S. aureus* under low pH conditions by helping bacteria to maintain their cytosolic pH.

## Discussion

In this study, we employed a Tn-Seq method to identify genes which are essential or detrimental for the growth of *S. aureus* in acidic pH. We used two large independently constructed transposon libraries for our experiments, and two stress conditions, pH 4.5 and pH 5.5. As expected, a greater number of genes were highlighted as being essential for growth at pH 4.5 compared to pH 5.5, with 31 and 5 genes, respectively, based on our analysis criteria. Hence, we primarily focused our work on the pH 4.5 genes and pathways. Differences in how the libraries were made, grown, or harvested resulted in approximately 50% of each gene set being unique to one library. But by combining the data from the two libraries and focusing only on the overlap, most of the genes identified in this manner could be confirmed, since reduced growth or survival at low pH was seen for mutants with individual genes inactivated.

A previously uncharacterized gene identified as essential for growth under acids stress conditions in our Tn-Seq experiment was *0846*. This gene was chosen for further study due to its low ratio when comparing the number of transposon insertions under the stress condition versus growth at neutral pH, indicative of being one of the most important genes under acid stress. This was further highlighted by the fact that this gene was also identified as one of only five genes identified as being essential for growth at pH 5.5. It is also interesting to note that 0846 has been identified as a virulence factor in several previous studies (37, 38), however, the actual cellular function of the protein was not known. 0846 is annotated as both a CPA transporter and a histidine permease. In a previous study deposited on a preprint server it has been shown that 0846, named NhaC3 in this study, has K^+^:H^+^ antiport activity at alkaline pH, and that its expression is upregulated at pH 9.0 (41). After performing amino acid uptake assays, it was however clear that 0846 is the main histidine transporter in *S. aureus*. To our knowledge, this is the first *in vivo* evidence of a histidine transporter in *S. aureus*. Our data analysing the growth of *S. aureus* in CDM at neutral pH with or without histidine indicates that the *0846::Tn* strain is still slightly sensitive to the presence of exogenous histidine. We hypothesise that *S. aureus* likely has another histidine transporter. An analysis of the *S. aureus* genome revealed gene *SAUSA300_0319* coding for the potential second histidine transporter (46). However, the radiolabelled histidine uptake assay, as well as the only slight sensitivity of the *0846::Tn* strain to exogenous histidine, demonstrates that 0846 is the main histidine transporter in *S. aureus*.

Suppressor strains of *0846::Tn* with increased acid tolerance all had mutations in *codY* or *relA* likely resulting in an activation of the stringent response. The stringent response in *S. aureus* is activated in response to a lack of amino acids, and results in increased expression of genes involved in amino acid uptake or biosynthesis. We hypothesise that increasing the expression of genes involved in histidine biosynthesis compensate for the lack of histidine transport in the *0846::Tn* mutant. Further confirmation of the importance of histidine for the survival of *S. aureus* under acid stress is our finding that the WT strain cannot grow in CDM at pH 4.3 without histidine, while it can when histidine is present. Consistent with this, the *0846::Tn* strain, lacking an active histidine transporter, could not grow in CDM pH 4.3 regardless of whether histidine was present or not in the medium.

*S. aureus* is capable of converting histidine into other compounds such as glutamate, where ammonia is produced as a by-product (47, reviewed in 48). Ammonia has been shown to be important for the acid stress response of *S. aureus*, either when produced via the urease enzyme or the ADI system (22, 23). We therefore initially hypothesised that the degradation of histidine via the histidine utilization (Hut) pathway (reviewed in 48), in which histidine is degraded to glutamate or glutamine and resulting in the production of two molecules of ammonia, was the reason for its importance. However, none of the *hut* genes, which are present in *S. aureus,* were identified as essential in the Tn-Seq assay. We also assessed *S. aureus* mutant strains with transposon insertions in *hutH* and *hutU,* coding for the enzymes catalysing the first two steps of this process, for their ability to grow under low pH conditions. Neither mutant display a growth defect on TSA pH 4.5 (data not shown), suggesting that degradation of histidine via the Hut pathway is likely not the reason why histidine is important during low pH growth conditions. Another degradation process of histidine that has been linked to the ability of bacteria to better grow in acid stress conditions is through a histidine decarboxylase system. In this pathway, histidine is decarboxylated to histamine and during this process a proton is consumed. While such systems have been shown to be important for the acid stress responses of lactic acid bacteria, and in particular *Lactobacillus sp.* (49), an analysis of the genome of *S. aureus* did not reveal any known bacterial histidine decarboxylase enzymes.

Histidine is an amino acid with a pKa of 6.0. This means that histidine is unique among amino acids for being able to act as a buffer within physiological pH ranges. As shown by our intracellular pH assays, the pH of WT *S. aureus* only decreases by 0.4 despite the extracellular pH decreasing from 7.3 to 4.5. This means that the cytoplasmic pH stays within the range that histidine is able to act as a buffer within. The *0846::Tn* strain however, was less able to buffer its intracellular pH compared to a WT strain. This effect was already seen at neutral pH but was even more pronounced under the low pH growth conditions. We hypothesise that the inability to take up histidine results in reduced intracellular concentrations of histidine, which means the mutant is less able to buffer its intracellular pH.

As validation of our Tn-Seq approach, we identified genes and pathways which have been previously associated with acidic stress in *S. aureus.* These included *vraG* and *dltB*, required for the maintenance of cell surface charge. The expression of *dltABCD* has been shown to be upregulated in response to acid shock (18), and recent work has shown that the GraXRS-VraFG five-component system may directly detect and respond to low pH (33, 34, 35). Our findings confirm the importance of modifying the cell surface charge to counteract acid stress. However, since most of the essential genes identified in our study code for proteins with known functions in cell wall assembly and maintenance, this indicates that besides cell surface charge, the cell wall as a whole has a more important role in helping bacteria to survive under acid stress than previously appreciated. Furthermore, some of the genes coding for proteins of still unknown cellular function and identified in this study as essential for growth in low pH, could potentially have a role in cell wall assembly.

We also identified several novel genes and pathways that are important for the growth of *S. aureus* in low pH. One of these was aerobic respiration, and in particular the QoxAB terminal oxidase. QoxAB accepts electrons from the electron transport chain to reduce oxygen to H_2_O. In the process, QoxAB transports protons out of the cell. Other bacterial proton pumps, in particular the F_0_F_1_-ATPase, have been shown to be important for the acidic stress responses of a range of bacteria, including *E. coli* (50), *S. enterica* (51, 52), and *Listeria monocytogenes* (53, 54, 55). However, it is unclear whether proton pumps act directly to buffer the intracellular pH, or that this movement of protons is required for ATP generation for use in other acidic stress responses. It is notable however, that *S. aureus* has a second terminal oxidase Cyd. The Cyd oxidase differs from QoxAB by not acting as a proton pump, which may explain why it was not identified as essential in our Tn-Seq experiment. This may support the hypothesis that the importance of QoxAB for growth in low pH is due to its proton pumping ability.

In conclusion, we have identified a wide range of pathways that are important for the growth of *S. aureus* under low pH. Some of these pathways have already been associated with the *staphylococcal* acidic stress response, such as the cell wall and cell surface charge, which validates our Tn-Seq method. Additionally, we identified several novel pathways, including aerobic respiration, and histidine transport via 0846.

## Materials and Methods

### Bacterial strains and growth conditions

Bacterial strains used in this study are listed in Table S5. *Escherichia coli* strains were grown in lysogeny broth (LB) or on LB agar plates and *Staphylococcus aureus* strains were grown on tryptic soy agar (TSA) plates or in tryptic soy broth (TSB). Where specified, the medium or plates were adjusted to the indicated pH with HCl prior to autoclaving. For the preparation of low pH agar plates, the bactoagar concentration was increased from the standard 15 % (w/v) to 30 % (w/v) as has been described previously (24). *S. aureus* strains were also grown in chemically defined medium (CDM), which was prepared as previously described (56, 57) or CDM lacking histidine. If required, the medium was supplemented with antibiotics as indicated in Table S5.

### Plasmid and bacterial strain construction

Several strains used in this study were derived from the Nebraska transposon mutant library (NTML) library (36). The transposon insertion site was confirmed by PCR and sequencing for all NTML strains used in this study. The *0846::Tn* transposon region from the original NTML strain NE967 (JE2 *0846::Tn*) was moved by phage transduction using phage Φ85 into fresh *S. aureus* JE2 or LAC* background strains, yielding strains JE2 *0846::Tn* transduced (ANG6197) and LAC* *0846::Tn* (ANG6049). For complementation analysis, plasmid pCL55-*0846* was constructed. The *SAUSA300_0846* gene including its native promoter region was amplified by PCR using primers 3443 (AAA<u>GAATTC</u>GAATTACCGATTACTGCAACCGAACGTGC) and 3444 AAA<u>GGATCC</u>GTCTCTAATAAATGAGTCATATTTTCACC). The resulting PCR product and plasmid pCL55 were digested with EcoRI and BamHI, ligated and initially recovered in *E. coli* strain CLG190 yielding strain CLG190 pCL55-*0846* (ANG6028). Plasmid pCL55-*0846* was subsequently isolated from *E. coli* and electroporated into the *S. aureus* strain RN4220, where it integrates into the *geh* gene locus, generating strain RN4220 pCL55-*0846* (ANG6069). Finally, this region was transduced with phage Φ85 into LAC* *0846::Tn* (ANG6197), resulting in the construction of the complementation strain LAC* *0846::Tn* pCL55-*0846* (ANG6076). Strain LAC* pCL55 (ANG3795) was used as control strain for some experiments and the empty plasmid pCL55 was also moved using phage Φ85 from *S. aureus* strain RN4220 pCL55 (ANG266) into strain LAC* *0846::Tn* (ANG6197) generating strain LAC* *0846::Tn* pCL55 (ANG6078). Strain LAC* *codY::Tn* (ANG6293) was constructed by moving the *codY::Tn* region from strain JE2 *codY::Tn* by phage Φ85 transduction into the LAC* strain background.

### Tn-Seq experiment

Two independent transposon mutant libraries were used in this study. Both libraries were constructed in the *S. aureus* USA300 TCH1516-derived MRSA strain TM283. The first library (library A) was a pool of around 600,000 colonies and was generated using a mix of six different transposons (containing 5 different outward facing promoters and one promoter less blunt transposon). Its construction was described previously (25, 58). The second library (library B) contained > 1 million colonies and was generated using only a single, promoter less transposon (blunt transposon). Construction of this library was also described previously (26). To identify genes essential during acid stress, the transposon mutant libraries were propagated in TSB medium (neutral pH of 7.3) or TSB medium adjusted with HCl to pH 5.5 or pH 4.5. A vial of the library was thawed on ice and used to inoculate 20 ml TSB medium supplemented with Erm 5 or 10 µg/ml to an OD_600_ of 0.1. This pre-culture was grown at 37°C with shaking for 1 h. This pre-culture was subsequently used to inoculate 25 ml (library A) or 100 ml with Erm 5 µg/ml (library B) of fresh TSB medium (pH 7.3), TSB pH 5.5 or TSB 4.5 medium to an OD_600_ of 0.00125. These cultures were grown at 37°C with shaking at 180 rpm until they reached an OD_600_ of around 1.4 (10 generations), bacteria from 12 ml culture were harvested by centrifugation, washed once with TSB and the cell pellet frozen at -20 °C for subsequent isolation of genomic DNA. The sample preparation for the sequencing analysis was performed as previously described (58). Briefly, the genomic DNA was digested with NotI, DNA fragments > 300 bp selectively precipitated with PEG8000, and then biotinylated adaptors ligated. Following this, the DNA was digested with MmeI, and Illumina adaptors with barcodes ligated. Samples were sequenced using an Illumina HiSeq platform and 100 base single end reads at the TUCF genomics core facility at TUFTS University, USA. The Illumina sequence reads for the TN-seq experiment using library A were deposited in the short read archive (SRA) at the National Center for Biotechnology Information (NCBI) under BioProject ID PRJNA998095. The Illumina sequence reads for the TN-seq experiment using library B were deposited as part of a previous study in the short read archive (SRA) under BioProject ID PRJNA544248 and accession number SRX5883253 (26).

### Tn-Seq data analysis

Analysis and mapping of the sequencing data was performed as previously described (25, 58) using the Tufts Galaxy Server. Briefly, reads were trimmed, split by Illumina barcode, and then further split by transposon barcode. Reads were then mapped to the USA300-TCH1516 genome and Hopcount tables generated. Statistical analysis was performed using the Mann-Whitney test to find significant differences in the number of reads per gene, and Benjamini-Hochberg test to correct the p-value for multiple hypothesis testing. Circular plots and Volcano plots were generated using R-scripts and circos. Further analysis of the Tn-Seq data was performed using filtering functions in Excel. The following parameters were used for filtering: Benjamini-Hochberg of ≤ 0.1 and only genes with ≥ 10 transposon insertions were considered. Genes were considered as essential under pH stress, when the ratio of the number of transposon insertions in the gene following growth under the low pH stress condition compared to the number of transposon insertions following growth in standard TSB (neutral pH) was ≤ 0.5 or as dispensable/detrimental genes when the ratio was ≥ 2. While the USA300-TCH1516 genome was used for the Tn-seq data analysis, USA300 FPR3757 locus tag numbers are shown in Table 1 to match up with the NTML strain annotations.

### Agar plate spotting assays

The indicated *S. aureus* strains were grown overnight in 3-5 ml TSB medium at 37 °C with shaking. The next day, bacteria from the equivalent of 1 ml culture with an OD_600_ of 5 were pelleted by centrifugation for 3 min at 17,000 xg and washed once with 1 ml phosphate buffered saline (PBS). Next, 10-fold dilutions down to a dilution of 10^-7^ were prepared in PBS. Five µl of the undiluted culture and each of the dilutions were spotted on standard TSA plates (pH 7.3) or TSA pH 4.5 plates acidified with HCl prior to autoclaving. Unless otherwise state, the experiments were done in three independent runs.

### Acid stress survival curves

The indicated strains were grown overnight in 3-5 ml TSB medium at 37 °C with shaking. Bacteria from 1 ml culture with an OD_600_ of 8 were collected by centrifugation at 17,000 xg for 3 min and washed 3 x with 1 ml TSB. The bacterial suspension was afterwards transferred to 20 ml TSB with a pH of 2.5 to yield an approximate OD_600_ of 0.4. Cultures were incubated at 37 °C with shaking at 180 RPM until the indicated time points. Colony forming units (CFU) were determined by removing 200 µl of bacterial culture and preparing 10-fold dilutions in PBS. 100 µl of selected dilutions were plated on TSA plates, and the plates were incubated at 37 °C for 16-18 h before colonies were counted. For the T = 0 h time point, culture aliquots were taken immediately after transfer to the low pH medium. The CFU count at T = 0 h was set to 100% for each strain and % survival at the subsequent time points calculated. The average and standard deviations of the % survival from three independent experiments were plotted.

### Bacterial growth curves in a plate reader

Growth curves were performed in various media as defined in the results section and figure legends. TSB or CDM was acidified to the indicated pH by the addition of HCl prior to autoclaving or filter sterilising. For some experiments, CDM without histidine was used. For the growth assays, bacteria were grown overnight in 3-5 ml TSB medium at 37°C with shaking. Bacteria from a 1 ml aliquot were pelleted by centrifugation for 3 min at 17,000 xg and washed once with 1 ml PBS buffer. The washed bacterial suspension was used to inoculate 1 ml fresh medium to a final OD_600_ of 0.05. Three technical repeats of 200 µl diluted culture were transferred to wells of a 96-well plate, and the plate was incubated at 37 °C with shaking at 500 rpm for 300 s every 10 minutes in a SPECTROstar^NANO^ plate reader. OD_600_ measurements were taken every 30 min for 24 h. The final OD_600_ values were calculated by averaging the three technical replicates and subtracting the OD_600_ readings of a blank well containing medium only. The averages and standard deviations of three independent experiments were calculated and plotted.

### Histidine uptake assays

Uptake assays measuring the incorporation of a radioactively labelled amino acid were performed as previously described (57). Briefly, cultures were grown overnight in CDM at 37 °C with shaking. Next day, the cultures were diluted to an OD_600_ of 0.05 into 50 ml fresh CDM and grown at 37 °C with shaking at 180 rpm to mid-log phase. Bacterial cells from 2 ml of culture were pelleted by centrifugation and washed once with 2 ml CDM lacking histidine and subsequently adjusted with CDM lacking histidine to an OD_600_ value of 1. An OD_600_ reading was taken again at this point, to determine the exact density of this culture suspension for normalization purposes. Next, to 450 µl of this washed bacterial culture, 2 µl of ^3^H radiolabelled L-histidine (histidine L-[ring-2,5-^3^H; ARC UK Limited, ART 0234) was added and the suspension mixed by swirling. Immediately afterwards (T=0 min time point), 100 µl culture was removed and filtered onto a nitrocellulose filter using a vacuum manifold and subsequently washed with 32 ml PBS. The remainder of the culture was incubated at room temperature (RT) and further 100 µl samples were taken and processed as described above at T=3, 6, and 9 min. The washed filters were subsequently removed and placed into 9 ml Filter-Count scintillation fluid (PerkinElmer). The radioactivity for each sample was measure in counts per minute (CPM) using a Wallac 1409 DSA liquid scintillation counter. CPMs were then normalised to the final OD_600_ values of the cultures used for these assays. Three independent experiments were performed, and average values and standard deviations calculated and plotted.

### Generation of LAC* *0846::Tn* suppressor strains with increased acid resistance

Multiple independent cultures of strain LAC* *0846::Tn* were grown overnight in TSB medium at 37°C. Next day, 50 µl of 10^-2^ to 10^-4^ dilutions were plated onto TSA pH 4.4 or 4.5 plates. Following 48 h incubation at 37°C, colonies obtained on the low pH TSA plates were re-streaked on TSA pH 4.4 or pH 4.5 and the plates incubated again for 48 h at 37 °C. Next, single colonies were picked, grown overnight in TSB supplemented with 10 µg/ml Erm and aliquots of these suppressor strain cultures stored frozen at -80°C. The increased acid resistance of the LAC* *0846::Tn* suppressor strains was subsequently confirmed using the agar plate spotting assay described above.

### Whole genome sequencing sample preparation and analysis

Genomic DNA was isolated using a previously described method (59, 60) and samples prepared for Illumina sequencing using an Illumina Nextera DNA kit. Samples were sequenced at the MRC London Institute for Medical Sciences in Hammersmith Hospital, using a Miseq machine and an Illumina MiSeq Reagent kit v2 (300 cycles) to generate 150 bp paired-end reads. For genome sequence analysis, a previously described protocol using the CLC workbench Genomics software (Qiagen) was used (61), and the reads were mapped to a custom *S. aureus* USA300 LAC* reference genome generated in a previous study, and for which the illumine reads have been deposited into the European Nucleotide Archive (ENA) under project number PRJEB14759 (24). Single nucleotide polymorphisms (SNPs) were determined based on at least 70% frequency, and strain background SNPs and areas of low coverage regions removed by comparing them to the SNPs and areas of low coverage also present in the genome sequence of the parental LAC\**0846::Tn* strain. Illumina sequence reads for the suppressor strains were deposited into the European Nucleotide Archive (ENA) under project number PRJEB61451.

### Intracellular pH measurements

Measurements of the intracellular pH were performed using the pHrodo^TM^ Red AM intracellular pH indicator dye (ThermoFisher, Cat. No. P35372) as described previously with some modifications (38). Overnight cultures of LAC* WT and LAC* *0846::Tn* were grown in TSB pH 7.3 or TSB pH 5.5 medium at 37°C with shaking. The following day, the cultures were diluted to a starting OD_600_ of 0.05 into 20 ml TSB pH 7.3 or TSB pH 4.5, respectively and grown to mid-log phase. Bacteria from the equivalent of 5 ml culture with an OD_600_ of 0.5 were collected by centrifugation for 20 min at 660 xg and subsequently washed 2 x with 5 ml 50 mM HEPES buffer pH 7.4. Next, following the manufacturer’s instructions, 20 µl of the PowerLoad concentrate was added to 2 ml of each culture and the bacteria stained with 5 µM pHrodo red dye final concentration for 30 min at RT in the dark. Following this incubation, 350 µl of the stained bacteria were collected by centrifugation for 3 min at 17,000 xg and suspended in 350 µl ml fresh 50 mM HEPES buffer pH 7.4. Two technical replicates of 150 µl were removed to a black-walled 96-well plate, before the fluorescence was measured on a TECAN plate reader with excitation / emission wavelengths of 560 / 590 nm at optimal gain. Measurements were taken every 5 min for 25 min, with shaking at 432 rpm for 120 s after each measurement. Intracellular pH values were determined by comparing the average of the two technical replicates of fluorescence measurements to a standard curve generated independently for each strain and pH growth condition to correct for differential dye uptake. For the generation of the standard curve, bacterial cultures were pre-grown to mid-log phase and stained with pHrodo Red dye as described above for the experimental sample. Following staining, four x 350 µl of culture per strain and condition were removed, bacteria collected by centrifugation for 3 min at 17,000 xg and resuspended in calibration buffers set to pH 4.5, pH 5.5, pH 6.5 or pH 7.5 and containing 10 µM nigericin and 10 µM valinomycin according to the manufacturer’s protocol (Intracellular pH Calibration Buffer Kit, ThermoFisher, Cat. No. P35379). The average of two technical replicates per calibration sample was plotted on a graph, and a semi-log linear regression used to calculate a line of best fit. This was used to convert fluorescence readings of the experimental sample into intracellular pH values. Three independent experiments were performed and averages and standard deviations of the intracellular pH at the T=10 min time point were plotted where the pH values fell within the ranges of the standard curve.

## Acknowledgments

This work was supported by the Wellcome trust grant 210671/Z/18/Z/WT to A.G. We thank Lisa Bowman, Freja CM Kirsebom and Sophie A Howard for their help with on of the transposon mutant library experiments.

