## Supplementary material for "Histidine and its uptake are essential for the growth of *Staphylococcus aureus* at low pH": Table S5 and Figures S1 - S3

**Supplemental Material**

**(5 Tables and 3 Figures)**

**Table S1. Tn-Seq data for Library A following growth at pH 4.5 compared to pH 7.3**

**Table S2. Tn-Seq data for Library B following growth at pH 4.5 compared to pH 7.3**

**Table S3. Tn-Seq data for Library A following growth at pH 5.5 compared to pH 7.3**

**Table S4. Tn-Seq data for Library B following growth at pH 5.5 compared to pH 7.3**

**Table S5. Bacterial strains used in this study.**

**Figure S1: Circular plots showing the transposon insertion density along the *S. aureus* genome at different pH growth conditions**

**Figure S2: Growth plate analysis of *S. aureus* mutant strains with transposon insertions in genes identified as essential for growth at pH 4.5.**

**Figure S3: Growth plate analysis of *S. aureus* mutant strains with transposon insertions in genes identified as dispensable for growth at pH 4.5.**

**Table S5:** Bacterial strains used in this study

| Unique ID | Strain name and resistance | Reference |
|---|---|---|
| | ***E. coli* strains** | |
| ANG243 | XL1-Blue pCL55; Amp 100 µg/ml | (1) |
| ANG1141 | CLG190 | Dana Boyd; (2) |
| ANG6028 | CLG190 pCL55-*0846*; Amp 100 µg/ml | This study |
| | ***S. aureus* strains** | |
| ANG3729 | TM283 (USA300-TCH1516 without pUSA300HOUMR) | (3, 4) |
| ANG2624 | JE2 (WT) | (5) |
| ANG4019 | JE2 *graS::Tn* (NE1756); Erm 10 µg/ml | (5) |
| ANG4008 | JE2 *vraG::Tn* (NE70); Erm 10 µg/ml | (5) |
| ANG4223 | Strain 923 (WT) | (6) |
| ANG4226 | Strain 923 Δ*vraS*; Cam 5 µg/ml | (6) |
| ANG4227 | Strain 923 Δ*vraR*; Cam 5 µg/ml | (6) |
| ANG6295 | JE2 *mprF::Tn* (NE1360); Erm 10 µg/ml | (5) |
| ANG5968 | JE2 *fmtA::Tn* (NE1022); Erm 10 µg/ml | (5) |
| ANG1575 | LAC* (WT) | (7) |
| ANG2395 | LAC* Δ*dltD* | (8) |
| ANG3969 | JE2 *SAUSA300_0482::Tn* (NE251); Erm 10 µg/ml | (5) |
| ANG3911 | JE2 *SAUSA300_0957::Tn* (NE1384); Erm 10 µg/ml | (5) |
| ANG5979 | JE2 *sagB::Tn* (NE1909); Erm 10 µg/ml | (5) |
| ANG3909 | JE2 *spdC::Tn* (NE1099); Erm 10 µg/ml | (5) |
| ANG5969 | JE2 *srrA::Tn* (NE1309); Erm 10 µg/ml | (5) |
| ANG3941 | JE2 *qoxB::Tn* (NE732); Erm 10 µg/ml | (5) |
| ANG5957 | JE2 *qoxA::Tn* (NE92); Erm 10 µg/ml | (5) |
| ANG5967 | JE2 *SAUSA300_0846::Tn* (NE967); Erm 10 µg/ml | (5) |
| ANG6058 | JE2 *SAUSA300_0846::Tn* (NE967); Erm 10 µg/ml (sequenced) | (5) |
| ANG6197 | JE2 *SAUSA300_0846::Tn* transduced; Erm 10 µg/ml | This study |
| ANG5970 | JE2 *SAUSA300_2389::Tn* (NE1400); Erm 10 µg/ml | (5) |
| ANG5963 | JE2 *SAUSA300_0429::Tn* (NE620); Erm 10 µg/ml | (5) |
| ANG5964 | JE2 *SAUSA300_0543::Tn* (NE802); Erm 10 µg/ml | (5) |
| ANG5959 | JE2 *SAUSA300_0481::Tn* (NE188); Erm 10 µg/ml | (5) |
| ANG5972 | JE2 *SAUSA300_1518::Tn* (NE1474); Erm 10 µg/ml | (5) |
| ANG5955 | JE2 *SAUSA300_1636::Tn* (NE22); Erm 10 µg/ml | (5) |
| ANG5966 | JE2 *SAUSA300_lepA::Tn* (NE865); Erm 10 µg/ml | (5) |
| ANG5978 | JE2 *SAUSA300_0759::Tn* (NE1891); Erm 10 µg/ml | (5) |
| ANG2631 | JE2 *noc::Tn* (NE486); Erm 10 µg/ml | (5) |

| | | |
|---|---|---|
| ANG4016 | JE2 *SAUSA300_2055::Tn* (NE1495); Erm 10 µg/ml | (5) |
| ANG5971 | JE2 *SAUSA300_1043::Tn* (NE1462); Erm 10 µg/ml | (5) |
| ANG5608 | JE2 *lytH::Tn* (NE1369); Erm 10 µg/ml | (5) |
| ANG6129 | JE2 *codY::Tn*; Erm 10 µg/ml | (5) |
| ANG6049 | LAC* *SAUSA300_0846::Tn*; Erm 10 µg/ml | This study |
| ANG6293 | LAC* *codY::Tn*; Erm 10 µg/ml | This study |
| ANG113 | RN4220 (WT) | (9) |
| ANG266 | RN4220 pCL55; Cam 7.5 µg/ml | (10) |
| ANG6069 | RN4220 pCL55-*0846;* Cam 7.5 µg/ml | This study |
| ANG3795 | LAC* pCL55, Cam 7.5 µg/ml | (11) |
| ANG6078 | LAC* *SAUSA300_0846::Tn* pCL55; Erm 10 µg/ml, Cam 7.5 µg/ml | This study |
| ANG6076 | LAC* *SAUSA300_0846::Tn* pCL55-*0846*; Erm 10 µg/ml, Cam 7.5 µg/m | This study |
| ANG6039 | JE2 *cpa1-1::Tn* (NE1504); Erm 10 µg/ml | (5) |
| ANG6040 | JE2 *cpa1-2::Tn* (NE366); Erm 10 µg/ml | (5) |
| ANG4556 | JE2 *cpa2::Tn* (NE308); Erm 10 µg/ml | (5) |
| ANG6037 | JE2 *nhaC1::Tn* (NE1470); Erm 10 µg/ml | (5) |
| ANG6038 | JE2 *nhaC2::Tn* (NE1214); Erm 10 µg/ml | (5) |
| ANG6080 | LAC* *SAUSA300_0846::Tn S-1;* Erm 10 µg/ml | This study |
| ANG6085 | LAC* *SAUSA300_0846::Tn S-2;* Erm 10 µg/ml | This study |
| ANG6087 | LAC* *SAUSA300_0846::Tn S-3;* Erm 10 µg/ml | This study |
| ANG6090 | LAC* *SAUSA300_0846::Tn S-4;* Erm 10 µg/ml | This study |
| ANG6099 | LAC* *SAUSA300_0846::Tn S-5;* Erm 10 µg/ml | This study |
| ANG6111 | LAC* *SAUSA300_0846::Tn S-6;* Erm 10 µg/ml | This study |
| ANG6113 | LAC* *SAUSA300_0846::Tn S-7;* Erm 10 µg/ml | This study |
| ANG6118 | LAC* *SAUSA300_0846::Tn S-8;* Erm 10 µg/ml | This study |

**Supplemental Figures:**

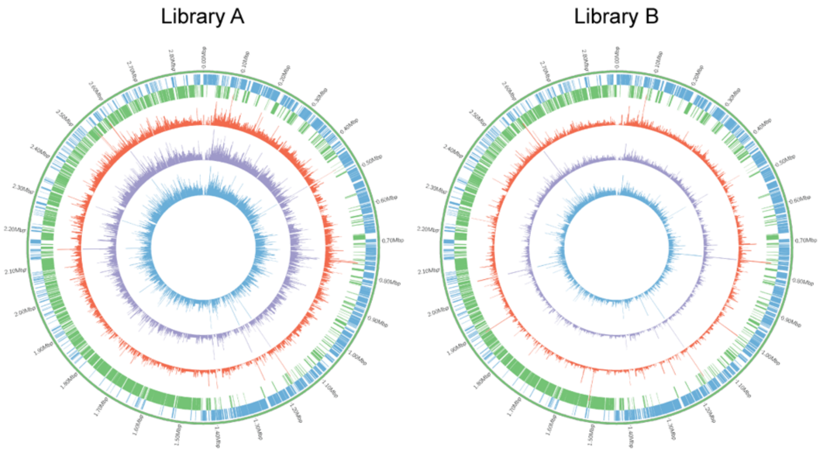


**Figure S1: Circular plots showing the transposon insertion density along the *S. aureus* genome at different pH growth conditions**. Circular plots for Tn-libraries A and B with the two outer rings depicting genes located on the (+) (blue) or (−) (green) strand in *S. aureus* strain. The inner three rings show the histograms of transposon insertions on a per gene basis after growth of the libraries in TSB pH 7.3 (red), pH 5.5 (purple), or pH 4.5 (blue) for 10 generations.

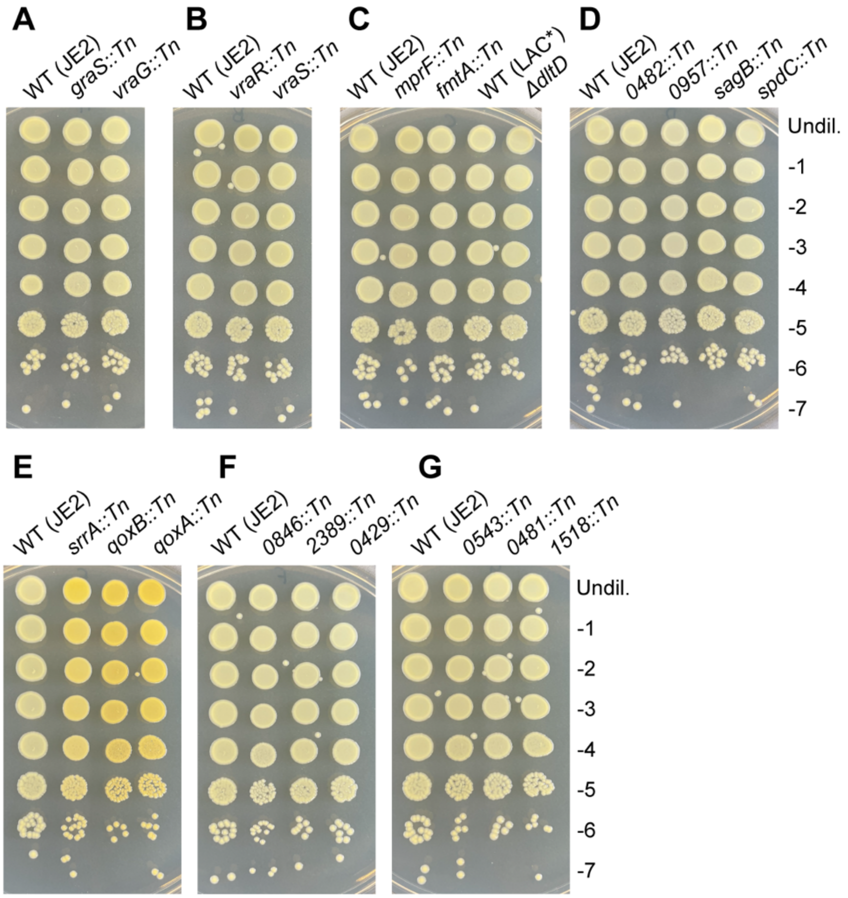


**Figure S2: Growth plate analysis of *S. aureus* mutant strains with transposon insertions in genes identified as essential for growth at pH 4.5.** (A-H) Bacterial growth on TSA pH 7.3 plates. Overnight cultures of the indicated WT and mutant strains were serially diluted and spotted on TSA pH 7.3 plates. Images were taken following 24 h incubation at 37 °C. Each image is a representative of three experiments.

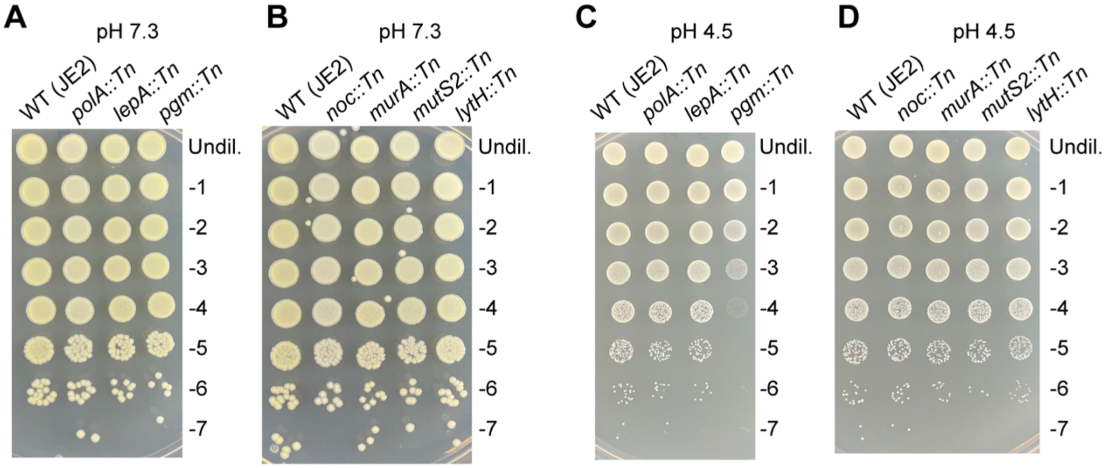


**Figure S3: Growth plate analysis of *S. aureus* mutant strains with transposon insertions in genes identified as dispensable for growth at pH 4.5.** (A-D) Bacterial growth on TSA plates. Overnight cultures of the indicated WT and mutant strains were serially diluted and spotted on (A-B) TSA pH 7.3 plates or (C-D) TSA pH 4.5 plates. Images were taken following 24 h incubation at 37 °C. Each image is a representative of three experiments.
